# Benchmark of force fields to characterize the intrinsically disordered R2-FUS-LC region

**DOI:** 10.1101/2022.12.20.521322

**Authors:** Maud Chan-Yao-Chong, Justin Chan, Hidetoshi Kono

## Abstract

Amyloid fibrils formations are involved in many neurodegenerative diseases such as Alzheimer’s disease, Parkinson disease, Amyotrophic Lateral Sclerosis (ALS) and others. The proteins associated with the formation of amyloid fibrils are Intrinsically Disordered Proteins (IDP) in the unbound state. Nevertheless, this type of proteins can self-aggregate and form cross-β amyloid fibrils structures at physiological condition.

Due to the flexibility of these IDPs, no single experimental approach could completely characterize this system, especially in the unbound state. All-atom molecular dynamics (MD) simulations could be used to study the conformational ensemble of IDPs. Unfortunately, force fields (FF) and water models (WM) were developed to simulate one structure of folded proteins. Recently, several FF/WM were improved to properly generate conformational ensembles of IDP. However, it is unknown if the force fields (FF) can properly reproduce the behavior of IDP and also self-aggregate in cross-β amyloid fibrils structures.

In this paper, we will focus of the R2 region of the FUS-LC domain (R2-FUS-LC region) which is an Intrinsically Disordered Region (IDR) of 16 residues in the unbound state but forms cross-β fibrils in the bound state. For the R2-FUS-LC region, we benchmarked thirteen commonly used FFs for studying IDPs. We show that CHARMM36m (updated in 2021) with mTIP3P water model performs the best to generate extended structures and cross-β amyloid fibril.

## INTRODUCTION

Proteins involved in amyloid fibril formation are intrinsically disordered and can self-aggregate to form cross-β structures^1^. Amyloidogenesis, formations or growth of amyloid fibrils, are involved in many pathologies including neurodegenerative diseases^2,3^ such as Alzheimer’s, Parkinson’s, type II diabetes, Amyotrophic Lateral Sclerosis (ALS)^2,4,5^ and others. However, there are many other transient structures and functions that are not well characterized^6–8^ such as its ability to form membrane-less compartment with droplet-like behavior, a characteristic of liquid–liquid phase separation (LLPS)^6,8–14^. At present, our understanding of the mechanism of amyloid fibrils aggregation and toxic intermediates^15^ is poor and this is one of the most reasons why various drug design strategies^16–18^ against neurodegenerative diseases face difficulties^19,20^.

ALS is rare neurodegenerative disease, with prevalence of 6 per 100 000 and incidence is around 2 per 100 000 population per year^21^. ALS is characterized by progressive muscular paralysis due to degenerative of motor neurons in the brain stem, spinal cord, primary motor cortex and corticospinal pathways^22^. In 50% of cases, death occurs within three years of the first clinical manifestation^23^. In ALS patients, amino acid mutations have been found in the Low-Complexity (LC) sequence regions of RNA granule forming proteins such as Fused in Sarcoma (FUS)^9,10,24–27^. Mutation in FUS-LC domain form irreversible amyloid fibril aggregation while the wild-type has reversible fibrils^9,27,28^.

The human FUS protein (526 residues), involved in mRNA splicing and transcription, can be decomposed into four domains: N-terminal LC domain, RGG-rich domain, RNA recognition motif, and C-terminal RG-rich domain (**Figure 1**).

**Figure 1.**
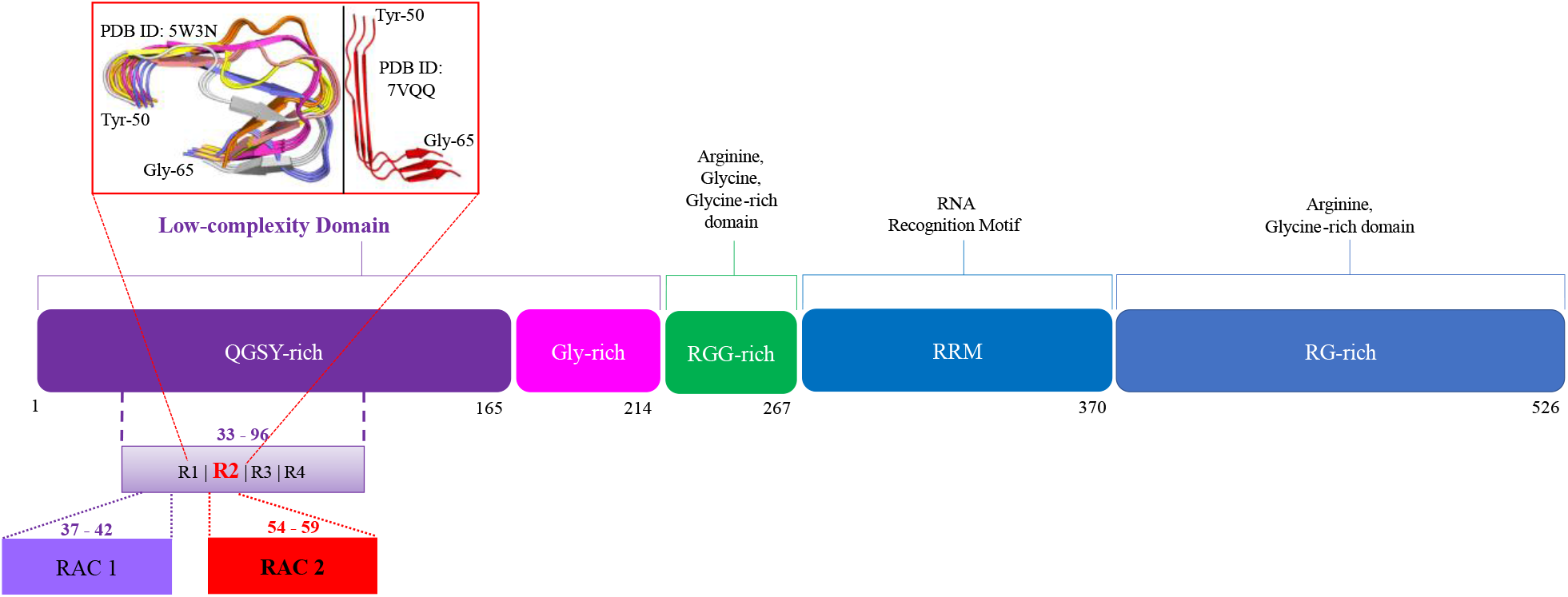
Domain organization of Fused in Sarcoma (FUS) protein which is constituted of 526-residue. FUS contains: N-terminal low-complexity sequence (LC) domain (residues 1–214) including a QGSY-rich prion-like domain (1–165, violet box) and a G-rich region (166–214, pink box); RGG-rich domain (215-267, green box); RNA-recognition motif (268-370, dark blue box); and C-terminal domain composed of RG repeat region (371-526, blue box). Inside on the red square, two different conformations of the R2-FUS-LC region is represented. On the left, six representatives of R2-FUS-LC region selected from the 20 models of the NMR structure (PDB ID: 5W3N^27^, “U-shaped”). On the right, the cryo-EM model from PDB ID: 7VQQ^29^, “L-shaped”.

The FUS N-terminal LC_1-214_ domain, under different conditions, is able to form different molecular assembles like LLPS, fibrils polymorphs^9,10,27–30^, reversible or irreversible cross-β amyloid fibrils^9,28,31,32^ while in the unbound state, the FUS-LC domain is intrinsically disordered.

In general, the sequence of the LC domain has little diversity and is composed of mostly polar or acidic amino acids as single amino acid repeats^33,34^ or repeats of the [S/G]Y[S/G] motif^9,28,35,36^. The full-length FUS-LC_1-214_ domain contains the FUS-LC-core_37-97_ implicated in amyloid fibril formation. The FUS-LC-core_37-97_ has four repeat motifs (R1-R2-R3-R4) containing multiple the [S/G]Y[S/G] motifs^28^(**Figure S1**, blue boxes). Within R1/R2, the tandem [S/G]Y[S/G] motifs have been implicated in the formation of Reversible Amyloid fibril Cores (RAC) (**Figure 1** and **Figure S1**)^28^. Lao *et al*. suggested that the S54/S61 and Y55 in R2 region of FUS-LC-core (R2-FUS-LC region) are involved in the stability of the RAC^37^ while Ding *et al*. showed that R2 is the most fibrilized repeat region^31^.

Although the two different structures of R2-FUS-LC have been experimentally solved^27–29,38^, any pathway or folding mechanism between IDR and amyloid fibril were solved. One possible solution is to perform all-atom Molecular Dynamics (MD) simulations to study IDRs. However, all atom MD studies for IDRs are not always consistent with experimental data^39,40^. It is believed that one of the main reasons is that the all atom force fields (FFs) such as AMBER, CHARMM^41,42^, OPLS-AA^43,44^ and GROMOS^45,46^ were developed for stable globular proteins. In contrast, IDPs/IDRs typically adopt multiple unstable conformations and nonpolar residues are often exposed to the solvent. Several research groups have modified FFs to better reproduce experimental IDP data^39,47– 52^ (**Table S2**, Sup. Info. Section Benchmark of FFs and water models (WMs)). Improvement of the FFs/WMs were brought by the modification of the protein-water interaction (Lennard-Jones parameters) at level of FF and WM, dihedral parameters modification or the optimization of grid-based energy corrections maps (CMAP potential). It is still unclear if these modified FFs are generalizable across all IDPs.

In this work, MD force fields widely used for IDPs are examined for the R2-FUS-LC_50-65_ region containing RAC2 (_54_SYSSYG_59_) (**Figure 1** and **Figure S1**) if they sample conformations consistent with the experimental results. We score the FFs based on three criteria: the compactness of the fibrils, the intra-peptide contacts in the folded cross-β state and the secondary structure propensity. Surprisingly, most FFs fail to reproduce the experimental data. Among them, our score suggests that the CHARMM36m2021 FF with the mTIP3P water model is recommended for studying R2-FUS-LC.

## METHODS

This section describes the details of the all-atom MD simulations and the software used for analyzing and comparing the MD simulation results with experimental data.

### PREPARATION OF THE INITIAL CONFORMATIONS FOR MOLECULAR DYNAMICS SIMULATIONS

Multiple 3D-structures of FUS-LC domain have been solved by X-ray crystallography^28,38^, electron crystallography^28,53^, cryo-Electron Microscopy (cryo-EM)^29^, and NMR^27^. The details of the structures are listed in the **Table S1**.

From the 20 NMR models (5W3N^27^), we truncated residues 50 to 65 (R2-FUS-LC region) of the first three chains (trimer) which form a U-shaped fibril-β. The 20 models were classified into 6 clusters using the GROMACS tool *gmx cluster*^54^ with a RMSD cutoff of 1.7 Å (computed over the Cα). We used a representative structure from each cluster as the initial conformation (**Figure 1**, red square). We performed independent MD simulations for the six representatives.

### MOLECULAR DYNAMICS SIMULATIONS

#### GENERAL CONDITIONS

We performed all-atom MD simulations using the GROMACS software version 2020.4^55,56^. The initial conformation of R2-FUS-LC (663 atoms) was solvated in a cubic box of 80×80×80 Å^3^. To be in the same conditions as a NMR experiment^31^, 42 sodium and 42 chloride ions were added to obtain 137mM NaCl (∼17,000 water molecules). The final systems contain 52000-68000 atoms.

### MOLECULAR DYNAMICS SIMULATIONS

We tested and compared thirteen force fields (FFs) and water models (WMs) recently developed for IDPs (**Table S2**, Sup. Info. Section Benchmark of FFs and WMs). From the six initial conformations, six independent MD simulations were carried out to equilibrate the systems for 2 ns in total. The nonbonded interactions were treated using the smooth particle mesh Ewald method^57^ for the coulomb terms and a cutoff distance of 12 Å or between 8 Å to 12 Å for the van der Waals potentials for AMBER or CHARMM FFs, respectively. The length of solute and water covalent bonds involved in hydrogen atoms was kept constant using the LINCS^58^ and SETTLE^59^ algorithms, respectively, allowing us to integrate the equations of motion with a 2 fs time step. After energy minimization, constant-pressure, and temperature (NPT) simulations were performed at 1 bar and 300 K for 1 ns to equilibrate the system. Temperature is controlled by the v-rescale thermostat and pressure by the Berendsen barostat^60,61^ with the time coupling constants τ_T_ = 0.1 ps and τ_P_ = 0.5 ps, respectively. A second equilibration run was performed with Nose-Hoover thermostat and Parrinello-Rahman barostat^62–64^ with the time coupling constants τ_T_ = 0.5 ps and τ_P_ = 2.5 ps for 1 ns.

For production, six 500 ns long MD simulations for each of thirteen force fields were conducted under the same conditions as that in the second equilibration, giving an accumulated trajectory of 3 μs. As shown from previous studies of IDPs^65^, running multiple MD simulations with a diverse set of the initial structures improves the quality of the IDP conformational ensemble. We discarded the first 100 ns and saved snapshots every 20 ps, finally obtaining ensembles of 120,000 structures for each FF.

### DATA ANALYSIS

Most of the trajectory analysis were performed with MDAnalysis, MDtraj, DSSP and some with our own scripts. All figures were plotted with Matplotlib (Python module).

### RADIUS OF GYRATION

The radius of gyration (Rg) of all 16 residues’ trimer and individual peptides were computed with MDAnalysis. The Rg distribution of the trimer (**Figure 2, Figure S2**) was then fitted to two Gaussians using the mixture gaussian model. To evaluate the FFs, we calculated Rg-Score_*FF*_ (refer to the unnormalized U/L-shaped Rg score) which was the inverted absolute Z-score_*FF*,*k*,*i*_of the experimental Rg’s (Rg_exp,k_) from the U-shaped (Rg_exp,U_:10.0Å, averaged over 20 models) and the L-shaped (Rg_exp,L_:14.4Å) conformations, which is given by:

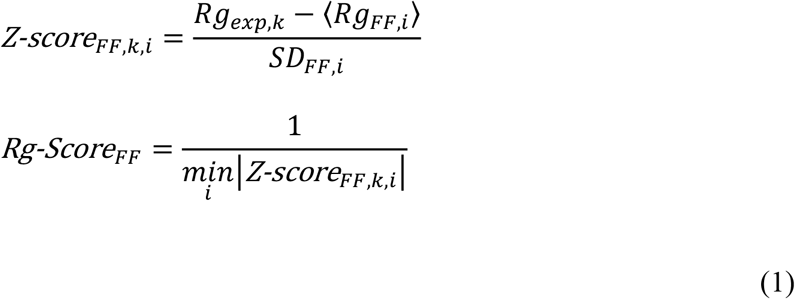

where <Rg_FF,i_> and SD_FF,i_ are the average and standard deviation, respectively, of the i’th Gaussian. For each FF, we took the largest inverted absolute Z-score_*FF*,*k*,*i*_ between the two Gaussians.

**Figure 2.**
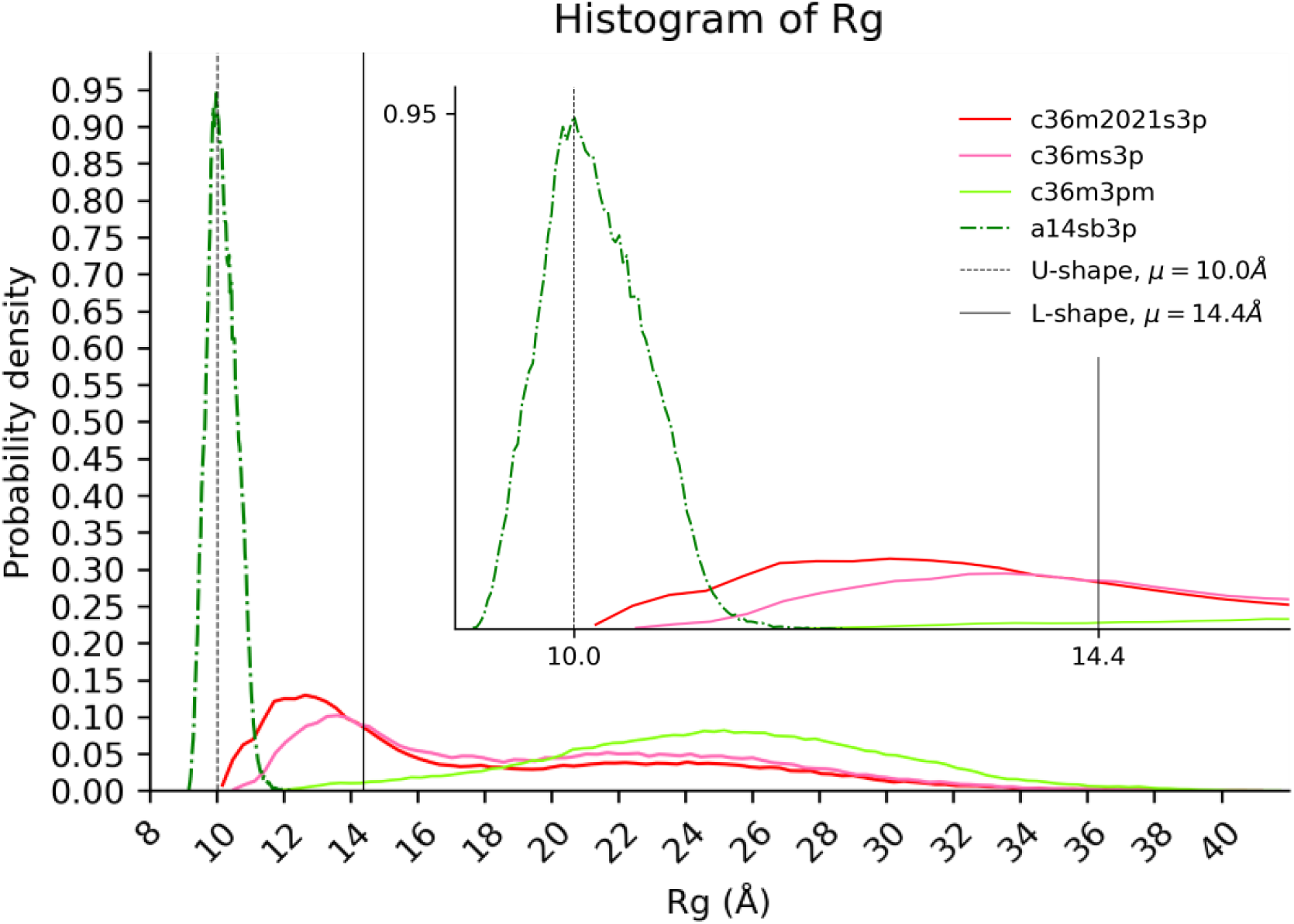
Distribution of the Rg for ensembles obtained with c36ms2021s3p (red), c36ms3p (pink), c36m3pm (light green) and a14sb3p (green).

The Rg of the individual peptides was compared to the predicted Rg_FL_ (10.8Å) from Flory’s polymer theory with parameters optimized for IDPs by Bernado and Blackledge^66^ and the Unfolded Rg score was calculated:

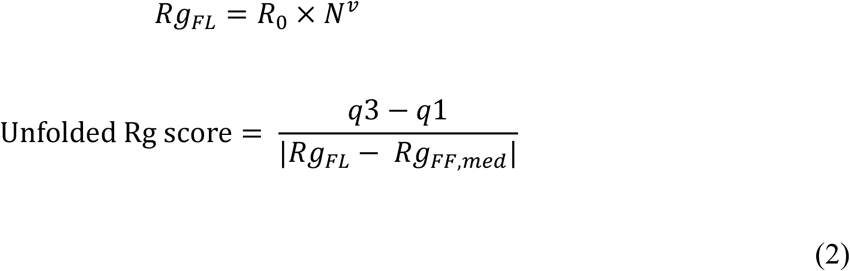

where R_0_ = 2.54 Å is the persistence length, ν = 0.522 is the exponential scaling factor, N is the number of residues, Rg_FF,med_ is the median Rg of the FF and q3 and q1 are the 75^th^ and 25^th^ percentile, respectively.

We linearly rescaled the Rg scores obtained from Eqs.(1) and (2) such that the top scoring FF was 1 and the bottom 0.0001. The final Rg score was computed in a way that product of the normalized U-shaped, L-shaped, and Unfolded Rg scores was calculated, and it was renormalized from 0.0001 to 1 (**Table 3**).

### CONTACT MAP ANALYSIS

Contact maps of heavy atoms within individual peptides were calculated using our own codes. Contact was counted when two atoms were within 5Å but excluded if they were in the flanking residues. The contact maps computed from the trajectories were compared to the reference contact map of the representative structure of U-shaped (**Figure 3**). For each FF simulation, an average of the trimer contact maps was calculated. We considered the contacts of which frequency was more than 1% in the trajectories.

**Figure 3.**
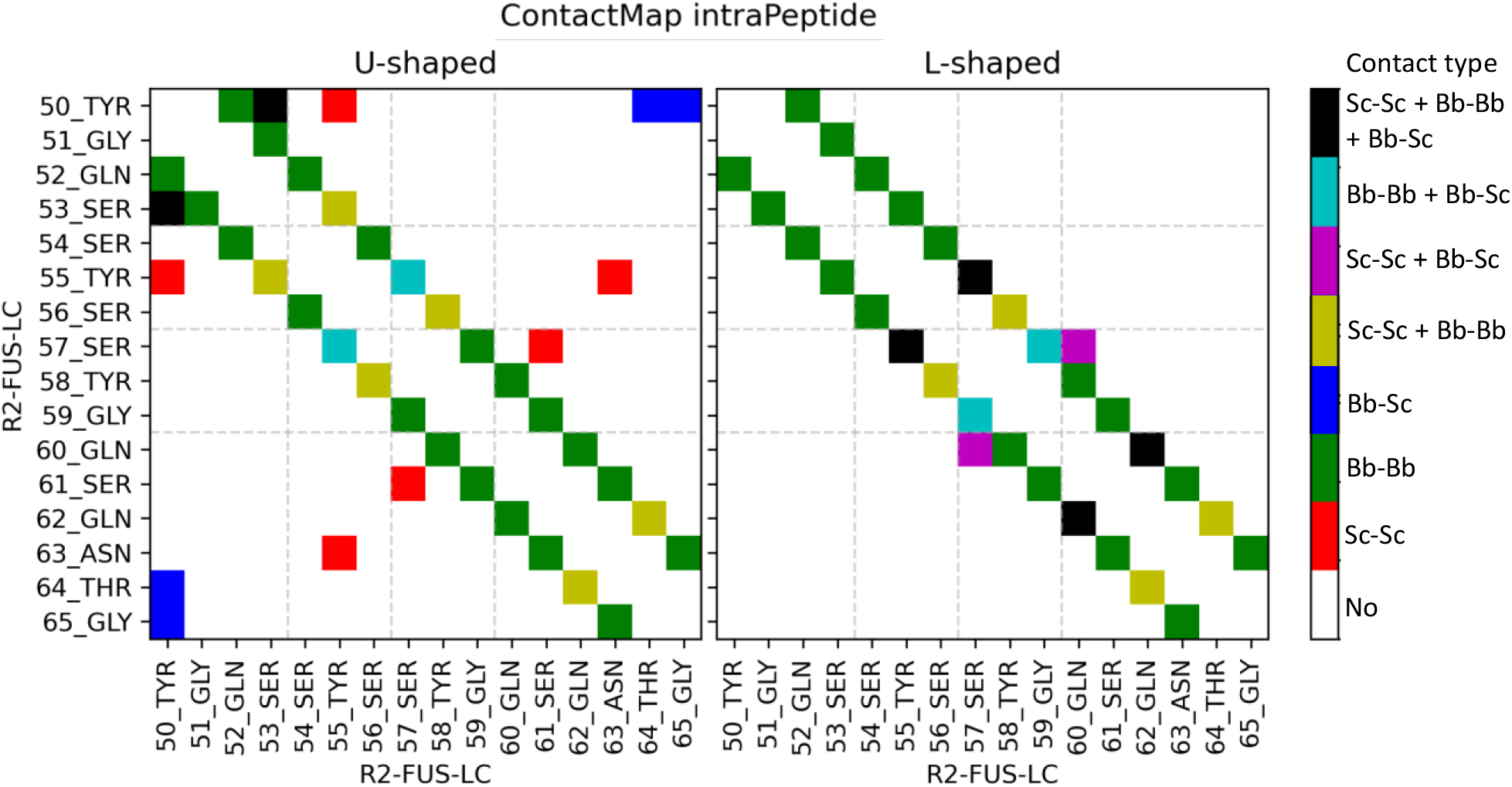
Contact Map intra-peptide for U-shaped (left) and L-shaped (right) conformations with 5 Å cutoff.

For each snapshot, contacts were classified into one of four groups [**Table 1**; True(T)/False(F) Positive(P)/Negative(N)] with respect to the reference contact map. Contact (or no contact) was considered positive (or negative).

**Table 1.** Confusion matrix.

|  |  | Contact in the Reference structure of U-shaped |  |
| --- | --- | --- | --- |
|  |  | Positive (P) | Negative (N) |
| Contact in the trajectory | Positive (PP) | True positive (TP) | False Positive (FP) |
|  | Negative (PN) | False Negative (FN) | True negative (TN) |

To measure the agreement of the contact maps, we computed the Matthews correlation coefficient (MCC):

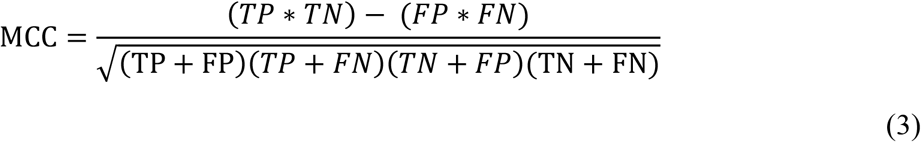

Note that the MCC score is between −1 and 1. In equation (3), true positive (TP), true negative (TN), false positive (FP) and false negative (FN) were calculated over all snapshots of the six replicas and average for the trimer (**Table S3**). MCC score of 1 shows perfect correlation with the reference, 0 with no correlation and −1 with perfectly negative correlation. We linearly rescale the top scoring as 1 and the bottom as 0.0001 (**Table 4**).

### SECONDARY STRUCTURE PROPENSITY

Secondary structure was determined using Dictionary of Protein Secondary Structure (DSSP)^67,68^. Since the assignment was changeable during the simulation, we calculated α-helix (H), β-strand (E) and coil propensities according to the occurrence. We used two measures for the comparison.

In the first measure, the assigned secondary structure from MD simulations was independently compared to the assigned secondary structure based on the U-shaped NMR (20 models) and L-shaped cryo-EM (single structure) (**Table S4**). We summed the log probability of the assigned secondary structure for the 16 residues in function of the U-shaped and L-shaped. We linearly rescaled the top SSP scoring as 1 and the bottom as 0.0001. We then combined the U-shaped and L-shaped to obtain the final SSP score (**Table 6**).

In the second measure, the secondary structure propensity for each 16 residues of the average of trimer from MD simulations was considered and they were plotted for α-helix and β-strand and compared to the U-shaped and L-shaped results (**Figure 4** and **Figure S2**).

**Figure 4.**
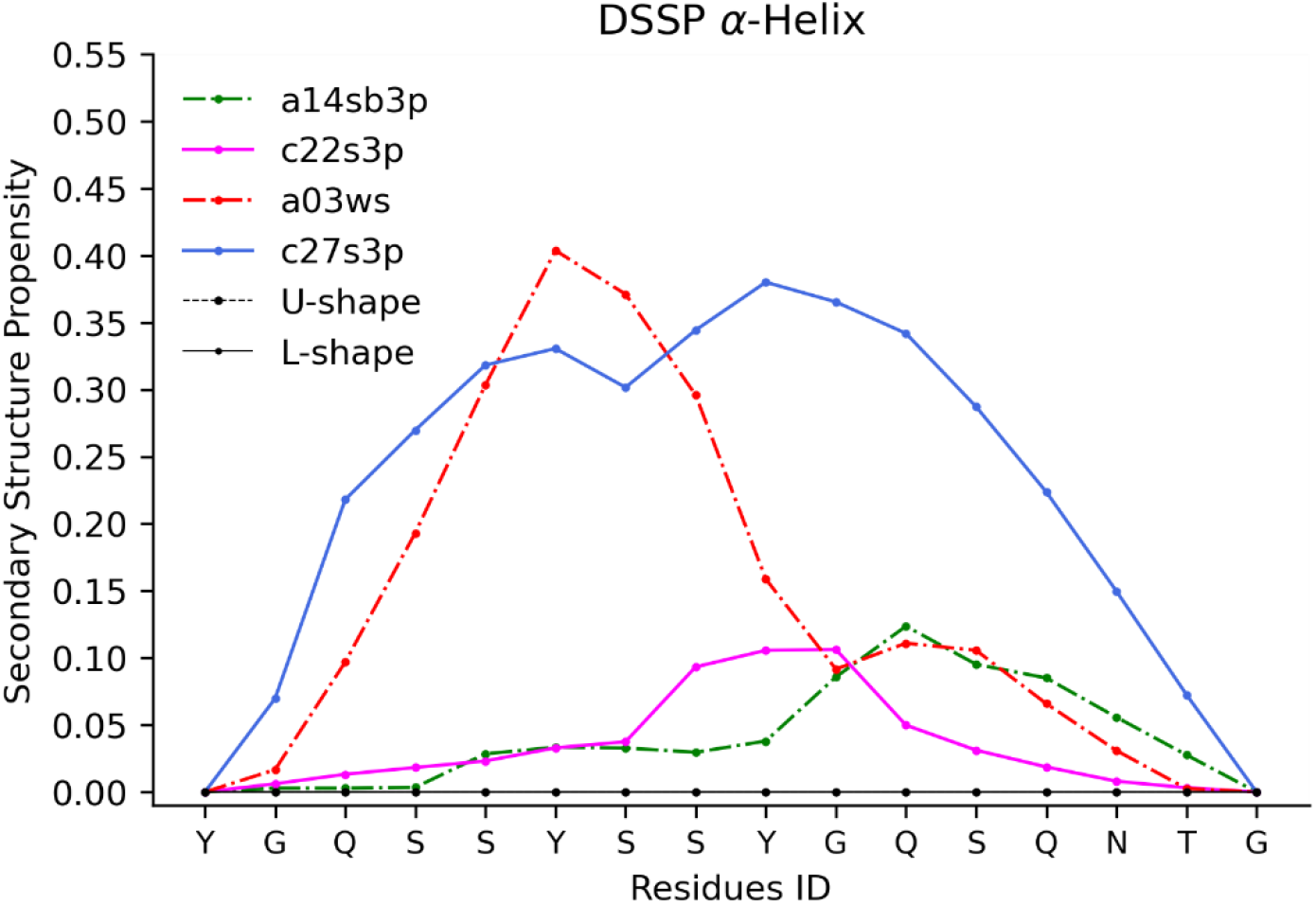
Probability Secondary Structure Propensity α-helices for FF forming helices (the values less than 0.1 were excluded).

### FINAL SCORING

Using the three distinct scorings (Rg score, SSP score and Contact map score), we calculated the final score for each FF by multiplying the scores and rescaling the product (from 1 to 0.0001) (**Table 2**).

**Table 2.** Summary of FF scores based on the three measures.

| Force Fields | Rg Score | SSP score | ContactMap score | Final Score | RANKING |
| --- | --- | --- | --- | --- | --- |
| **c36m2021s3p\*** | 1 | 0.65613 | 0.77657 | 5.10E-01 | 1 |
| **a99sb4pew\*** | 0.39845 | 1 | 0.64988 | 2.59E-01 | 2 |
| **c36ms3p\*** | 0.8692 | 0.31996 | 0.85621 | 2.38E-01 | 3 |
| **a19sbopc\*** | 0.30691 | 0.61874 | 0.7161 | 1.36E-01 | 4 |
| **a99disp\*** | 0.21986 | 0.51499 | 0.543 | 6.15E-02 | 5 |
| **a99sbildn4pd\*** | 0.19783 | 0.45716 | 0.47616 | 4.31E-02 | 6 |
| **a99sbCufix3p\*** | 0.07391 | 0.7345 | 0.34292 | 1.86E-02 | 7 |
| **c22s3p<sup>\*</sup>** | 0.05156 | 0.4001 | 0.53323 | 1.10E-02 | 8 |
| **a03ws<sup>#</sup>** | 0.02029 | 0.10312 | 0.1012 | 2.12E-04 | 9 |
| **a14sb3p<sup>#</sup>** | 0.00001 | 0.76161 | 0.40067 | 3.05E-06 | 10 |
| **c36m3pm<sup>#</sup>** | 0.00029 | 0.0032 | 0.9946 | 9.23E-07 | 11 |
| **c36m2021s3pm<sup>#</sup>** | 0.04438 | 0.00001 | 1 | 4.44E-07 | 12 |
| **c27s3p<sup>#</sup>** | 0.0313 | 0.00001 | 0.00001 | 3.13E-12 | 13 |

## RESULTS

We have carried out six, 500 ns-long MD simulations with each of the thirteen FFs (**Table S2**). We observed conformational transitions between the NMR conformation (the initial conformation) and extended conformations during the simulations for at least some of the FFs (**Figure S3**). Therefore, we decided to evaluate FF with 3μs-long trajectory in total for each FF.

To evaluate FF against the reversible amyloid fibril R2-FUS-LC region (wild-type trimer, each containing 16 residues), we used three measures: radius of gyration (Rg), secondary structure propensity and intra-peptide contact map. Rg can measure the global compactness/extension of the trimer’s peptides and the individual peptides. Secondary structure propensity and intra-peptide contact map both concern the local contact details of the R2-FUS-LC region. Based on the combined score based on these three measures (**Table 2, Final Score**), we ranked the thirteen FFs (see **Table 2**).

Based on the final score (**Table 2**), the thirteen FFs can be separated into three distinct groups: ^*^: top ranking group,•: middle ranking group, ^**#**^: bottom ranking group according to the order of scores.

FFs in the “top” group have medium (0.3-0.7) to high (>0.7) scores to the reference data for all measures. FFs in the “bottom” group, c27s3p and a03ws gave low scores (<0.3) in all the three measures. a14sb3p FF got relatively good scores for SSP and Contact Map scores, but low score for Rg. c36m3pm FF performed best for ContactMap score, but worst for SSP. FFs in “middle” ranking group tend to have low scores for at least one of the three measures but have medium agreement for the remaining. Details of the three measures will be explained in the following sections.

### .1. Global information of R2-FUS-LC region

The Rg score shows the ability of the FFs to sample the folded cross-β and unfolded states of the R2-FUS-LC region. The reference data for the folded cross-β structure state consists of two different conformations (“U-shaped”, PDB: 5W3N^27^ and “L-shaped”, PDB: 7VQQ^29^, **Figure 1**) of the R2-FUS-LC region forming a cross-β structure. Twenty U-shaped models with different loop conformations were solved by NMR with an average Rg of 10.0Å (measured as trimer of R2-FUS-LC region) and the less compact L-shaped conformation (trimer of R2-FUS-LC region Rg: 14.4Å) was solved by cryo-EM.

The cross-β structure is reversible^9,28^, implying that the R2-FUS-LC region can also adopt unfolded conformation. Intrinsically, structure in the unfolded state is not uniquely determined. As measure for structures in the unfolded state, we used a Flory’s random coil polymer model with optimized parameters for IDPs^66^ to estimate Rg of structure in the unfolded state.

The distributions of Rg for snapshots of the simulation data are shown in **Figure 2** and **Figure S2**. To evaluate how common/often the FFs can sample conformation close to the reference structures in terms of Rg, we fit Rg distributions with two Gaussian distributions. The distance to the reference Rg was computed as the absolute number of standard deviations from the mean(s) (Z-score_*FF*,*k*,*i*_) of the Gaussian(s). The lower Z-score_*FF*,*k*,*i*_ was chosen, inverted and normalized by linearly scaling them from 0.00001 to 1.0 (**Table 3**). The final Rg score was calculated by multiplying the three normalized scores and rescaling them linearly in a similar manner as Rg scores for individual three conformations (**Table 3**).

**Table 3.** Normalized Rg scores for the thirteen FFs.

| Force fields | Rg Score (Rank) |  |  | Final Rg Score<sup>a</sup> |
| --- | --- | --- | --- | --- |
|  | U-shaped | L-shaped | Unfolded |  |
| **c36m2021s3p\*** | 0.05654 (8) | 0.26227 (3) | 0.04817 (3) | 1.00000 |
| **c36ms3p\*** | 0.02194 (11) | 1.00000 (1) | 0.02830 (4) | 0.86920 |
| **a99sb4pew\*** | 1.00000 (1) | 0.43082 (2) | 0.00066 (12) | 0.39845 |
| **a19sbopc\*** | 0.11252 (4) | 0.17544 (6) | 0.01111 (5) | 0.30691 |
| **a99disp\*** | 0.09730 (5) | 0.16614 (7) | 0.00972 (7) | 0.21986 |
| **a99sbildn4pd\*** | 0.06440 (7) | 0.20090 (5) | 0.01092 (6) | 0.19783 |
| **a99sbCufix3p<sup>•</sup>** | 0.14855 (3) | 0.21916 (4) | 0.00162 (11) | 0.07391 |
| **c22s3p<sup>•</sup>** | 0.04650 (9) | 0.11735 (9) | 0.00675 (8) | 0.05156 |
| **c36m2021s3pm<sup>•</sup>** | 0.00035 (12) | 0.08977 (12) | 1.00000 (1) | 0.04438 |
| **c27s3p<sup>#</sup>** | 0.07513 (6) | 0.13992 (8) | 0.00213 (10) | 0.03130 |
| **a03ws<sup>#</sup>** | 0.03703 (10) | 0.10192 (10) | 0.00384 (9) | 0.02029 |
| **c36m3pm<sup>#</sup>** | 0.00001 (13) | 0.08983 (11) | 0.22404 (2) | 0.00029 |
| **a14sb3p<sup>#</sup>** | 0.91433 (2) | 0.00001 (13) | 0.00001 (13) | 0.00001 |

<sup>a</sup>Obtained by multiplication of the 3 scores and renormalizing them to 0.00001-1.0. FFs are sorted by the final Rg score in descending order.

The thirteen FFs can be separated into three groups. Comparing the ranking of the final score (**Table 2**) and the Rg final score only (**Table 3**), the top four FFs (c36m2021s3p, a99sb4pew, a19sbopc and c36ms3p) are the same although the order is slightly different.

Looking at **Table 3**, we noticed two types of FFs for the four top FFs. One type is c36m2021s3p and a19sbopc whose performance were uniform for three reference structures. The other is c36ms3p and a99sb4pew which have a bias for either flexible or compact conformations. This bias in sampling was also observed in a99sb4pew, a99sbCufix3p and a14sb3p for the U-shaped conformation, and c36m2021s3pm, c36m3pm for the unfolded conformation. However, their final Rg rank correlates with the strength of their bias as the scoring scheme penalizes an FF that fits well to one specific conformation but performs poorly for the others. Such FFs are the two worst performing FFs, c36m3pm and a14sb3p, which show opposite behaviors. c36m3pm sampled mostly unfolded conformations (**Figure 2**) while a14sb3p was biased towards the initial U-shaped conformation (**Figure 2**).

CHARMM FFs tend to generate more extended conformations than AMBER FFs (**Figure S2**), except for a03ws. AMBER FFs showed a bias towards the initial structure of the simulations, the folded cross-β amyloid fibril.

The Rg measures global compactness of conformation but is not proper to evaluate the detailed structures. For evaluating sampled structures in more detail, we introduced measures, contact map and secondary structure propensity (SSP), that evaluate the intra-peptide interactions.

### .2. Intra-peptide contacts of the R2-FUS-LC region

The U-shaped and L-shaped conformations contain 20 and 15 intra-peptide contacts, respectively (**Figure 3**). In the L-shaped conformation, no contact was observed between residues N and ≤N+5 (medium distance contacts) within 5 Å cutoff. In the U-shaped conformation, medium distance contacts are found between Tyr_50_-Tyr_55_, Tyr_50_-Thr_64_, Tyr_50_-Gly_65_, Tyr_55_-Asn_63_ and Ser_57_-Ser_61_, which can be used for FF evaluation.

We calculated the average intra-peptide contacts for the trimer using sampled conformations. The MCC score is used to compare the contacts from the FFs and the U-shaped conformation (**Table 4**). MCC is 1 (or −1) when two contact maps are perfectly correlated (or anti-correlated) and 0 if they have no correlation. MCC scoring penalizes false predictions, therefore FFs that have the most native contacts (TP>14.30) like a99sb4pew, c27s3p and a14sb3p (Table S3), were not necessarily the top scoring FFs (**Table 4**).

**Table 4.** Matthews Correlation Coefficient (MCC) of the FFs intra-peptide contact maps with respect to the U-shaped conformation’s contact map.

| Force Fields | Matthews Correlation Coefficient Score | Norm. Contact Map Score |
| --- | --- | --- |
| **c36m2021s3pm\*** | 0.70418 | 1.00000 |
| **c36m3pm#** | 0.70323 | 0.99460 |
| **c36ms3p\*** | 0.67892 | 0.85621 |
| **c36m2021s3p\*** | 0.66494 | 0.77657 |
| **a19sbopc\*** | 0.65431 | 0.71610 |
| **a99sb4pew\*** | 0.64268 | 0.64988 |
| **a99disp\*** | 0.62391 | 0.54300 |
| **c22s3p\*** | 0.62219 | 0.53323 |
| **a99sbildn4pd\*** | 0.61217 | 0.47616 |
| **a14sb3p<sup>#</sup>** | 0.59891 | 0.40067 |
| **a99sbCufix3p\*** | 0.58877 | 0.34292 |
| **a03ws<sup>#</sup>** | 0.54631 | 0.10120 |
| **c27s3p<sup>#</sup>** | 0.52854 | 0.00001 |

There are some surprising results in the contact map score ranking (**Table 4**).The c36m2021s3pm and c36m3pm FFs were top ranked in the contact map score although they were classified as a bottom performer, in terms of the final score (**Table 2**). c36m2021s3p, which was best in terms of the final score, was only ranked 4th. Furthermore, a99sbCufix3p was ranked in the bottom group though it was ranked in the middle in terms of the final score.

Looking at the confusion matrices of the four FFs in more detail (**Table 5** and **Table S3**), we have a few observations. First, c36m2021s3pm, c36m3pm and c36m2021s3p, which were best to the Unfolded Rg score, all preferred more extended conformations which were suggested by <20% of residues being in contact. This is in contrast with a99sbCufix3p which preferred more compact conformations (23.16% of residues in contact).

**Table 5.** Confusion Matrices.

| c36m2021s3p |  | Ref |  | Total Pred. |
| --- | --- | --- | --- | --- |
| Pred | T | 13.86 | 5.13 | 18.99 |
|  | F | 5.18 | 75.82 | 81.01 |

| a99sbCufix3p |  | Ref |  | Total Pred. |
| --- | --- | --- | --- | --- |
| Pred | T | 14.17 | 8.99 | 23.16 |
|  | F | 4.88 | 71.96 | 76.84 |

| c36m2021s3pm |  | Ref |  | Total Pred. |
| --- | --- | --- | --- | --- |
| Pred | T | 13.75 | 3.52 | 17.27 |
|  | F | 5.3 | 77.43 | 82.73 |

| c36m3pm |  | Ref |  | Total Pred. |
| --- | --- | --- | --- | --- |
| Pred | T | 13.75 | 3.57 | 17.32 |
|  | F | 5.29 | 77.38 | 82.67 |

Although extended conformations sampled with these FFs had fewer contacts, most of the native contacts were indeed observed in the sampled conformations. On the other hand, a99sbCufix3p had more contacts, but most were non-native contacts which appeared in the FP (**Table 5**).

Except for the three FFs, the remaining FFs ranking (**Table 4**) were similar to that in the final score similarly (**Table 2**). As mentioned earlier, the L-shaped conformation has no medium distance contact but it did form β-strand secondary structures which could not evaluated with the contact maps. We thus used the Secondary Structure Propensity (SSP) of these FFs and show the results in the next section.

### .3. Secondary Structure Propensity (SSP) of R2-FUS-LC region

In order to evaluate the FFs in terms of the SSP of the R2-FUS-LC region, we employed the log probability of observing the experimental (U-shaped and L-shaped) secondary structures given the SSP distributions obtained from the simulations.

In **Table 6**, we observe that the SSP score is higher for a14sb3p and a99sbCufix3p and lower for c36ms3p and c36m2021s3pm compared with their final rank. The FFs that are ranked higher in terms of the SSP score tend to produce more compact conformation, resulting in yielding the Rg values closer to those of the U/L-shaped conformations (**Table 3**). On the other hand, the FFs that are ranked lower tend to prefer a more extended conformation. These results indicate the importance of secondary structure consideration in the FF evaluation.

**Table 6.** Secondary Structure Propensity score to the U-shaped and L-shaped conformations.

| Force Fields | SSP Score U-shaped | SSP Score L-shaped | SSP Score |
| --- | --- | --- | --- |
| **a99sb4pew\*** | 1 | 1 | 1 |
| **a14sb3p<sup>#</sup>** | 0.852963 | 0.892893 | 0.761607 |
| **a99sbCufix3p<sup>\*</sup>** | 0.823343 | 0.892089 | 0.734498 |
| **c36m2021s3p\*** | 0.787453 | 0.833228 | 0.656131 |
| **a19sbopc\*** | 0.725248 | 0.853141 | 0.618743 |
| **a99disp<sup>\*</sup>** | 0.708013 | 0.72737 | 0.514992 |
| **a99sbildn4pd<sup>\*</sup>** | 0.677607 | 0.674657 | 0.457158 |
| **c22s3p<sup>\*</sup>** | 0.630039 | 0.635035 | 0.400103 |
| **c36ms3p\*** | 0.564177 | 0.567119 | 0.319962 |
| **a03ws<sup>#</sup>** | 0.330989 | 0.311533 | 0.103123 |
| **c36m3pm<sup>#</sup>** | 0.044558 | 0.071552 | 0.003198 |
| **c36m2021s3pm<sup>\*</sup>** | 0.00001 | 0.057822 | 0.00001 |
| **c27s3p<sup>#</sup>** | 0.011461 | 0.00001 | 0.00001 |

The a03ws and c27s3p FFs which are ranked at the bottom in terms of the SSP scoring have a strong preference for forming α-helices around the RAC2 motif (**Figure 4**). However, this region are β-strands in both the L-shaped^29^ and U-shaped^27^ structures. The NMR chemical shift data also shows that there are no α-helices throughout the R2-FUS-LC region^27,28,31,35,69^.

## DISCUSSION

We observed that AMBER FFs, a14sb3p and a99sbCufix3p tend to produce more compact conformations while the CHARMM FFs, c36ms3p and c36m2021s3pm, do more extended conformations. This trend seems to be true for other CHARMM and AMBER FFs.

CHARMM FFs systematically prefer more extended conformations (**Figure S2**). Four (all c36m FFs) of the six CHARMM FFs were top ranked in the terms of Rg score to unfolded conformations but ranked lower than 8 for either or both Rg scores to the U/L-shaped conformations (**Table 3**). This is further supported by the SSP score where CHARMM FFs are mostly ranked at the bottom.

While the AMBER FFs prefer more compact conformations (**Figure S2**) where six of the seven AMBER FFs are within the top 7 in the U-shaped Rg score (except a03ws) and three (a99sbCufix3p, a99sb4pew, a14sb3p) of the seven FFs are ranked at the bottom of the unfolded Rg score. AMBER FFs are biased towards compact structures, but the contact map analysis shows they have more contacts but the increase in contact is due to non-native contacts (**Table 5**).

However, the AMBER FFs perform well in the SSP score, indicating that they can reproduce the native secondary structures. Therefore, their poor ranking in the contact map score is due to non-native “medium distance” contacts.

Lao *et al*. used AMBER99SB-ILDN (with TIP3P water model) to simulate a longer region of FUS including the R2-FUS-LC region^37^. Comparing with their contact maps, we observe similar contact patterns in this study. Furthermore, the α-helix SSP of ∼5% in the previous work is similar to this study. However, they observed much higher β-strand SSP than we observe (**Figure S4**), probably due to the length of the peptide: 60 residues in the previous study while 16 residues in this study.

Studies on Amyloid-β proteins showed that different FFs give different conformational ensemble, but some are in agreement with the experimental fibril aggregation^52,70,71^. Similarly to our results, Pedersen *et al*. found ff19SB/OPC (equivalent to our name: a19sbopc) produces more compact conformations and forms more β-strand than a99SB-disp/TIP4P-disp (a99disp)^70^. For cross-β peptides, Samantray *et al*. demonstrated CHARMM36m gave promising result to sample and to give conformational ensemble with random coil and β-strand structures^71^, which is consistent with our results. Moreover, we find that our results are consistent to Piana *et al*. where they tested three and four-site water models with AMBER and CHARMM FFs^52^. When using TIP3P (3-site) water model, CHARMM FFs are more flexible than AMBER FFs. Using 4-site water models, AMBER FFs become a lot more flexible. This is consistent with what we observe except for a99sb4pew (**Figure S2**).

In more general, Robustelli *et al*., evaluated AMBER and CHARMM FFs against multiple globular, hybrid and IDPs^39^. Their ranking is similar in that our final score ranking except for C36m (c36ms3p) perform better than a99SB-*disp* (a99disp) and a99SB-ILDN/TIP4P-D (a99sbildn4pd). Looking at the SSPs, C22*/TIP3P (c22s3p) and a03ws (a03ws) overestimated the α-helix propensity just as we observed in our system (**Figure 4**).

## CONCLUSIONS

This study shows that force fields developed for globular proteins and modified for intrinsically disordered proteins, with their water models adapted, generate conformational ensemble essentially different to each other. We demonstrate here that it is important to combine multiple measures for a well-rounded evaluation of the FFs. This is due to the intrinsic flexibility of this system where the peptide can form β-strand fibrils and be in random coiled conformations.

Our benchmark of thirteen force fields from CHARMM and AMBER families indicates that c36m2021s3p is best balanced in terms of the three measures used for FF evaluation. This FF can generate conformational ensemble which contains both extended and cross-β amyloid conformations for the short region R2-FUS-LC domain and showed a good agreement in terms of the SSP and contact map. Some top FFs are biased to the initial structure, which is observed more for AMBER FFs. It should be noted that the combination of c36m2021and mTIP3P water model is computationally less inexpensive than those of top-ranked AMBER FFs with four-site water model.

## AUTHOR INFORMATION

### Author Contributions

The manuscript was written through contributions of all authors. All authors have given approval to the final version of the manuscript.

## ACKNOWLEDGMENT

MD simulations were performed using JAEA resources from QST institute and local resources.

## ABBREVIATIONS

ALS: Amyotrophic Lateral Sclerosis;
FF: force field;
FUS: Fused in Sarcoma;
FUS-LC domain: Low-Complexity domain of FUS;
IDP: Intrinsically Disordered Proteins;
IDR: Intrinsically Disordered Region;
LC: Low Complexity;
MD: molecular dynamics;
TDP43: Tar DNA-binding protein 43;
Rg: radius of gyration;
hnRNPA1: Heterogeneous nuclear ribonucleoprotein A1;
RNA: Ribonucleic Acid.

