## Supplementary Materials for "Benchmark of force fields to characterize the intrinsically disordered R2-FUS-LC region"

**Benchmark of Force fields (FF) and Water models (WM)**

In the late 20th century, all-atom protein FFs were developed to simulate the conformational dynamics of folded proteins (AMBER99[1], CHARMM22[2] with the three-site water model, SPC/E[3] or sTIP3P[4]), which generally have their nonpolar residues buried in their core and protected from solvent. This characteristic is observed only for globular and folded protein but not for IDPs / IDRs.

That is why, recently, several FF / WM were improved to properly generate conformational ensembles of IDPs / IDRs, in which nonpolar residues are frequently exposed to solvent while being suitable for globular proteins.

One approach is to modify the protein-water interaction at level of water model:

This is generally achieved by using a four-site water model, such as TIP4P[4] or OPC water model[5], which better accounts for the electric properties of water molecules, and by accentuating the depth of the solute-solvent Lennard-Jones (LJ) potentials to better solvate nonpolar residues.

TIP4P water model reparametrized with Ewald techniques provide the new water model TIP4P-Ew in 2004[6].

The second improvement was reported by Best et al., who proposed to rescale by a factor $\gamma$ = 1.1 the LJ parameters εOi between the water oxygen of model TIP4P/2005 and the protein atoms of AMBER-03w FF[7]. This modified water-solute interactions was denoted in the literature as TIP4P/2005s. The combination of AMBER-03w with TIP4P/2005s is named A03ws.

Another improvement, to counter the underestimation of water dispersion interaction with some polar amino acid, is to increase the Lennard-Jones dispersion coefficient $C_6$ = 900 kcal mol$^{-1}$ Å$^{6}$, for TIP4P/2005 and called the new water model TIP4P-D[8].

The first improvement from standard TIP3P (sTIP3P)[4] to CHARMM-TIP3P (mTIP3P)[2] was to add the $\varepsilon_H$ = -0.046 kcal $mol^{-1}$ and the $\sigma_H$ = 0.040 nm of the water hydrogen and the non-bonded interaction, and the $\varepsilon_{OH}$ = -0.0836 kcal $mol^{-1}$ and the $\sigma_{OH}$ = 0.177 nm between the water hydrogen and the water oxygen.

One improvement concerns the CHARMM FF by increase the Lennard-Jones well depth parameter $\varepsilon_H$ = -0.10 kcal $mol^{-1}$ of the water hydrogen of the specific mTIP3P water model, this water model named CHARMM-TIP3Pm (mTIP3Pm)[9].

The second approach is to modify this protein-water interaction at level of FF:

Another improvement concerns the AMBER FF. Yoo and Aksimentiev optimized the AMBER99SB-ILDN-Phi FF with their pair-specific Lennard-Jones parameters describing amine nitrogen-carboxylate oxygen interactions and also aliphatic carbon-carbon pairs thanks to experimental data of osmotic pressure from amino acid in solution, named this new FF AMBER99SB-ILDN-phi-CUFIX (A99Cufix3p)[10].

http://bionano.physics.illinois.edu/CUFIX

The third approach is to modify the dihedral parameters: This parameter modifies the energy of the system as a function of rotation around bonds. Since the flexibility of the biomolecule is correlated to the bond rotations, these corrections can affect the sampling of the conformational ensemble of the studied biomolecule.

The improvement for the FF was first focused on the backbone $\varphi$ and $\psi$ dihedral potentials to better reproduce the two-dimensional Ramachandran probability distributions like for

AMBER03w[11], CHARMM22*[12], AMBER14SB[13], AMBER19SB[14] FFs. AMBER19SB is the updated FF of AMBER99SB and AMBER14SB.

Another optimization improved sides chain torsion potentials to refine side-chain rotamers to improve the helix propensities, calling '-ILDN' extension of some FFs, like AMBER99SB-ILDN[15,16].

From this idea, a fourth approach is to optimize grid-based energy corrections maps (CMAP potential):

MacKerell et al. optimized CHARMM22 with correction maps based on cross-term energy functions of φ and ψ angles to better account for their correlations (CMAP) based on experimental NMR observables for backbone. This new FF is called CHARMM27[2,17].

MacKerell et al. goes even further by updating this CMAP potentials to correct the φ and ψ probability distributions for intrinsically disordered proteins, optimizing the side-chain dihedral potentials from IDP's NMR data[18], and updating from CAMP strategy and divided Cα into 3 different groups in function of the residue type. These modifications conduct to update CHARMM27 to CHARMM36 and to get CHARMM36m9.

Every year, some small updates for CHARMM36m for proteins-ions-lipids were adjusted. The last update used was in February 2021 where the NBFIX terms have been changed between chloride and sodium ions interaction.

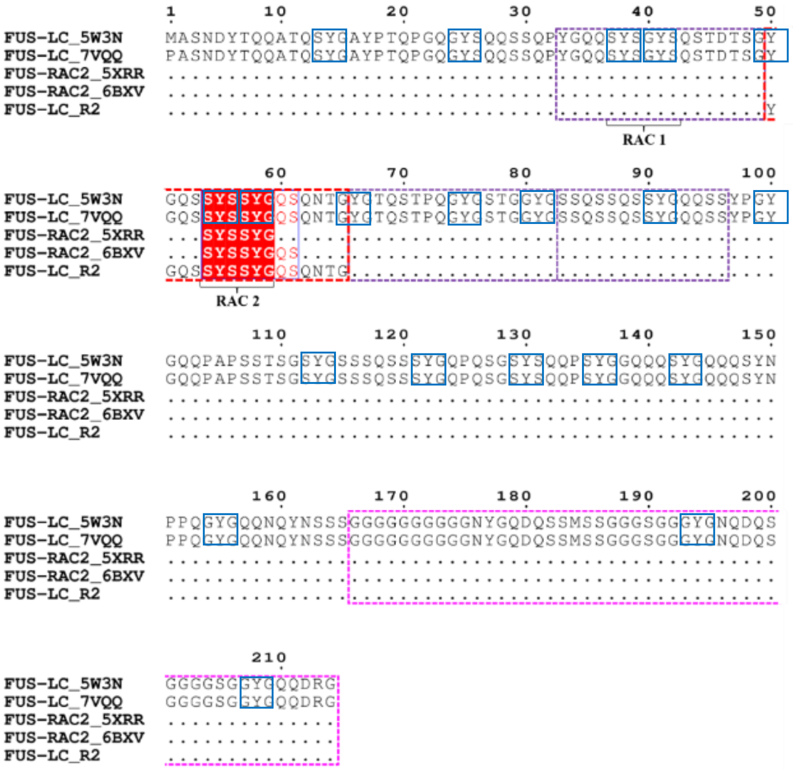

**Figure S1.** Sequence of the full-length FUS-LC$_{1\text{-}214}$ domain. Multiple sequence alignment of the human full-length Low-Complexity (LC) domain of FUS protein used by Murray *et al.* (FUS_5W3N, PDB ID: 5W3N). The amino acid numbering is based on the human protein sequence (UNIPROT ID: P35637). The 20 blue boxes correspond to the motif of three residues [S/G]Y[S/G]. The red box represents the R2-FUS-LC domain containing the reversible amyloid cores (RAC2) residues highlighted in red. The three violet boxes indicate the repeat region R1 (containing RAC1), R3 and R4, respectively. The pink box indicates the Gly-rich domain.

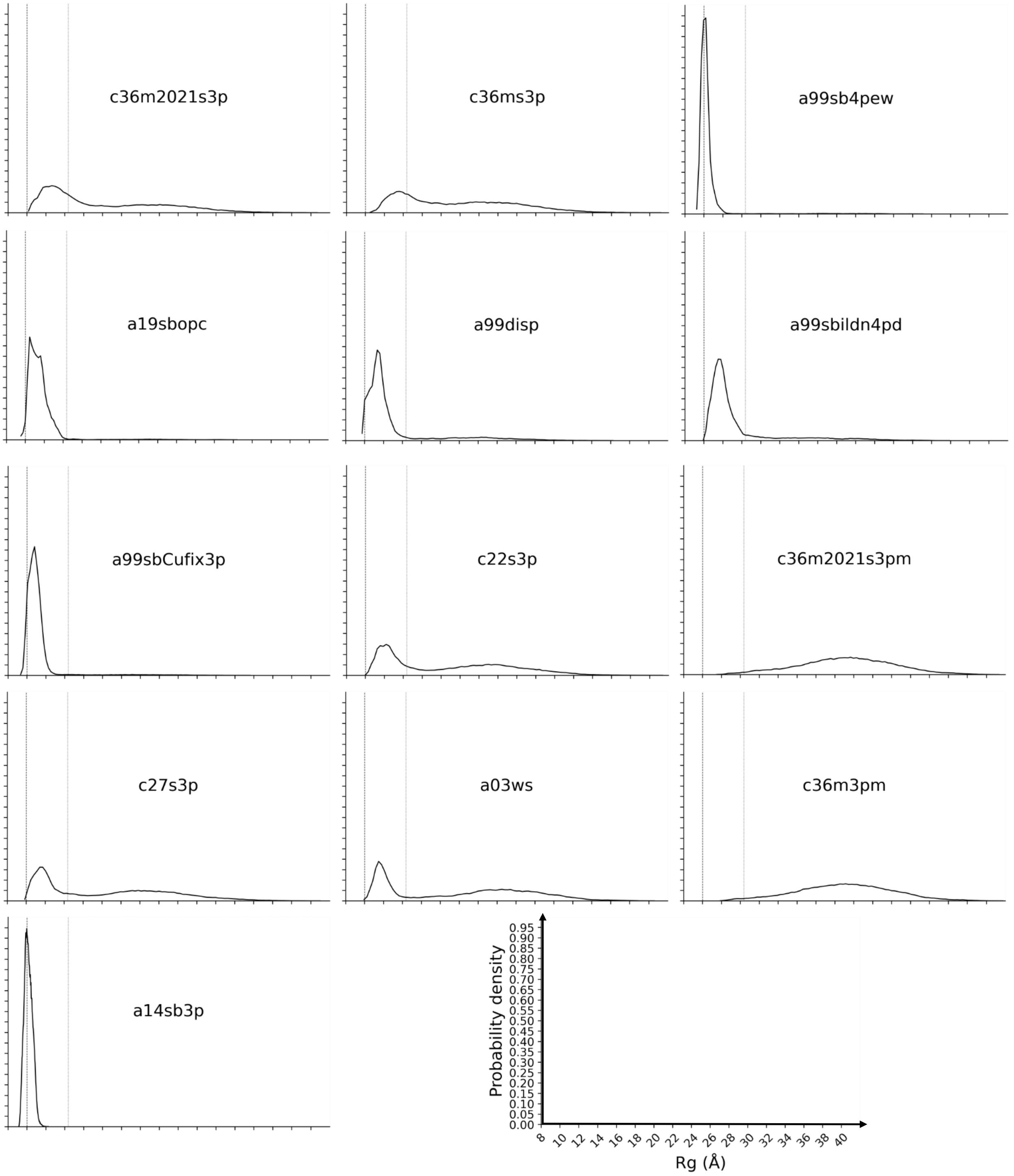


**Figure S2.** Distribution of the Rg for the (A) AMBER and the (B) CHARMM FFs. Vertical dash and dot lines correspond to the average of the Rg for U-shape (10.0 Å) and L-shape (14.4 Å), respectively.

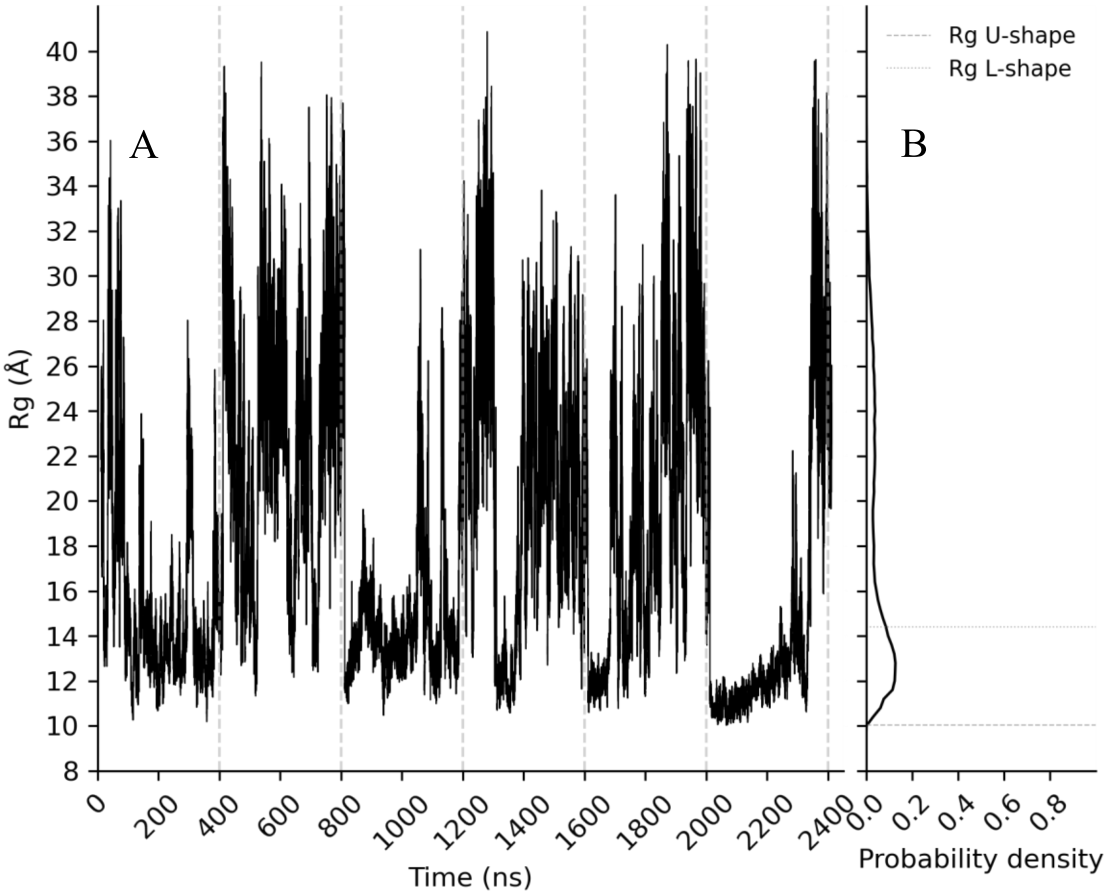


**Figure S3.** Rg in function of trajectory times (A) and distribution of the Rg (B) for c36m2021s3p FF.

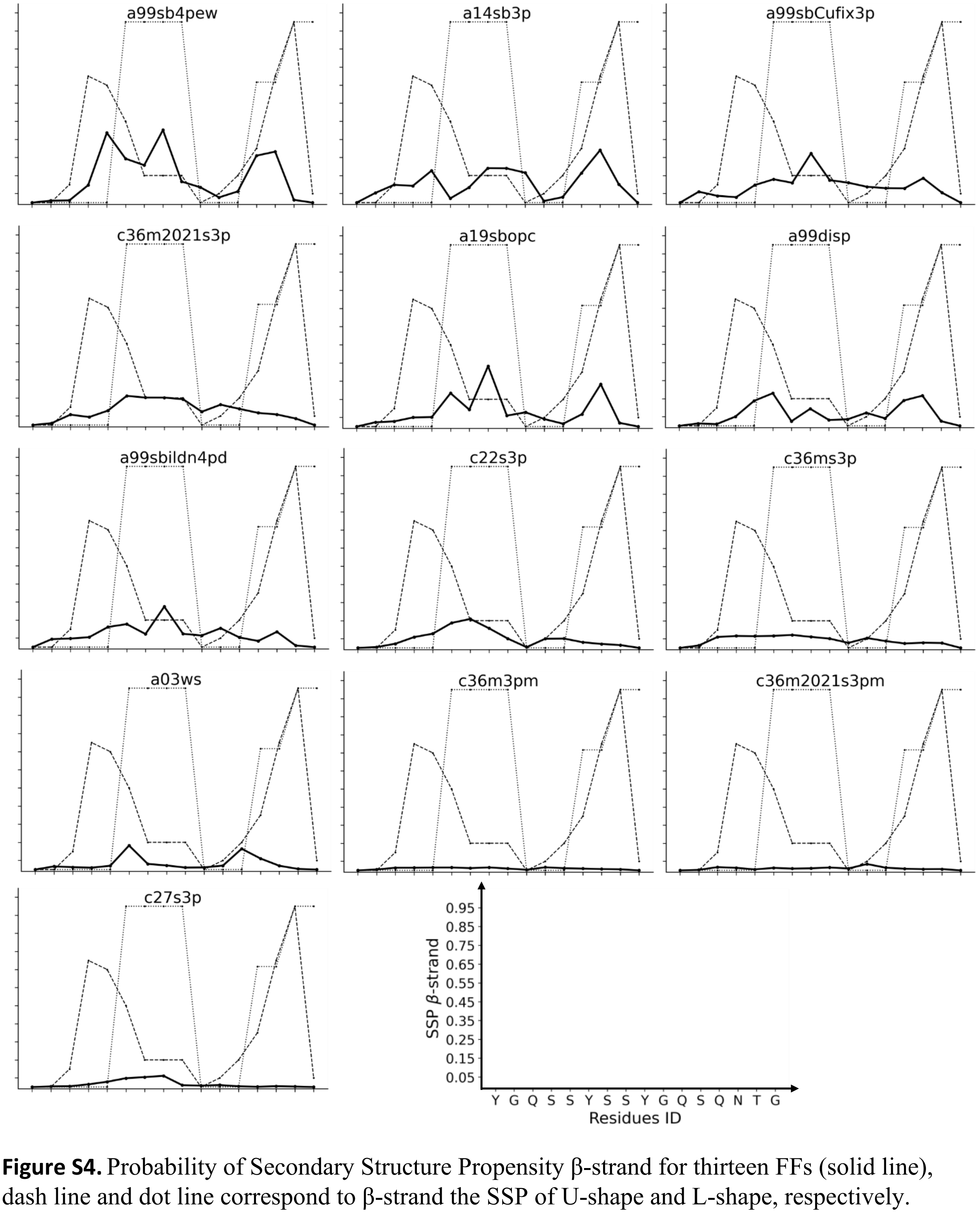


**Figure S4.** Probability of Secondary Structure Propensity β-strand for thirteen FFs (solid line), dash line and dot line correspond to β-strand the SSP of U-shape and L-shape, respectively.

**Table S1.** List of PDB structures from FUS-LC domain or short region of this domain. *: PDB ID with R2-FUS-LC region solved or partially solved.

| PDB ID | Year | Method | Purified Protein residues | Visible residues | Publication |
|---|---|---|---|---|---|
| **5W3N*** | **2017** | **NMR** | **2 - 214** | **37 – 97 (20 models)** | [19] |
| 5XSG | 2018 | Electron Crystallography (0.73 Å) | 37 – 42 | 37-42 | [20] |
| **5XRR*** | 2018 | X-ray (1.5 Å) | 54 - 59 | **54 - 59** | [20] |
| 6BWZ | 2018 | X-ray (1.1 Å) | 37-42 | 37-42 | [21] |
| **6BXV*** | 2018 | X-ray (1.1 Å) | 54 - 61 | **54 - 61** | [21] |
| 6KJ1 | 2019 | Electron Crystallography (0.65 Å) | 37-42 | 37-42 | [22] |
| 6KJ2 | 2019 | Electron Crystallography (0.67 Å) | 37-42 | 37-42 | [22] |
| 6KJ3 | 2019 | Electron Crystallography (0.60 Å) | 37-42 | 37-42 | [22] |
| 6KJ4 | 2019 | Electron Crystallography (0.65 Å) | 37-42 | 37-42 | [22] |
| **7VQQ*** | **2022** | **Cryo-EM (2.90 Å)** | **2-214** | **34-124** | [23] |

**Table S2.** Description of FFs and WMs.

| FF (Year) | Modifications | WM (Year) | Modifications | Abbreviation |
|---|---|---|---|---|
| AMBER99SB (2006)[24] | | TIP4P-Ew (2004)[6] | Protein-water interaction | A99sb4pew |
| AMBER03w (2010)[11] | Dihedral parameters (backbone) | TIP4P/2005s (2014)[7] | Protein-water interaction | A03ws |
| AMBER99SB-ILDN (2010)[15] | Protein-water interaction | TIP4P-D (2015)[8] | Protein-water interaction | A99sbildn4pd |
| AMBER14SB (2015)[13] | Dihedral parameters (Backbone) | TIP3P (1983)[4] | | A14sb3p |
| AMBER99SB-ILDN-phi-Cufix (2016)[10] | Protein-water interaction (Nonbonded fix strategy NBFIX) + Dihedral parameters (sidechain) | TIP3P (1983)[4] | | A99Cufix3p |
| AMBER19SB (2020)[14] | Dihedral parameters to improve helical propensities of A14SB (Backbone profiles for all 20 amino acids) | OPC (2014)[5] | Not parameterized against a specific atomistic WM for IDP | A19sbopc |
| AMBER99SB-Disp (2018)[25] | Optimized a99SB-ILDN by changing the protein backbone hydrogen bonds interaction | A99SB-disp water[25] | Optimized TIP4P-D by increasing the C6 dispersion term to avoid the over collapse the helical propensity | A99disp |
| CHARMM27 (2004)[2,17] | CMAP potential | mTIP3P (1998)[2,4] | Protein-water interaction ($\varepsilon/\sigma_H$ and $\varepsilon/\sigma_{OH}$) | C27s3p |
| CHARMM22* (2011)[2,12] | Dihedral parameters (Backbone and sidechain) | mTIP3P (1998)[2,4] | Protein-water interaction ($\varepsilon/\sigma_H$ and $\varepsilon/\sigma_{OH}$) | C22s3p |
| CHARMM36m (2017)[9] | CMAP potential (based on NMR data from IDPs) | mTIP3P (1998)[2,4] | Protein-water interaction ($\varepsilon/\sigma_H$ and $\varepsilon/\sigma_{OH}$) | C36ms3p |
| CHARMM36m (2017)[9] | CMAP potential (based on NMR data from IDPs) | mTIP3Pm (2016)[9] | Protein-water interaction ($\varepsilon_H$) for IDP | C36m3pm |
| CHARMM36m2021 (2021)[9] | CMAP potential (based on NMR data from IDPs) | mTIP3P (1998)[2,4] | Protein-water interaction ($\varepsilon/\sigma_H$ and $\varepsilon/\sigma_{OH}$) | C36m2021s3p |
| CHARMM36m2021 (2021)[9] | CMAP potential (based on NMR data from IDPs) | mTIP3Pm (2016)[9] | Protein-water interaction ($\varepsilon_H$) for IDP | C36m2021s3pm |

**Table S3.** Confusion matrix of the 13 FFs.

| c36m2021s3pm• | Ref_T | Ref_F |
|---|---|---|
| **Pred_T** | 13,75 | 3,52 |
| **Pred_F** | 5,30 | 77,43 |

| c36m3pm# | Ref_T | Ref_F |
|---|---|---|
| **Pred_T** | 13,75 | 3,57 |
| **Pred_F** | 5,29 | 77,38 |

| c36ms3p* | Ref_T | Ref_F |
|---|---|---|
| **Pred_T** | 13,86 | 4,61 |
| **Pred_F** | 5,19 | 76,34 |

| c36m2021s3p* | Ref_T | Ref_F |
|---|---|---|
| **Pred_T** | 13,86 | 5,13 |
| **Pred_F** | 5,18 | 75,82 |

| a19sbopc* | Ref_T | Ref_F |
|---|---|---|
| **Pred_T** | 14,02 | 6,16 |
| **Pred_F** | 4,71 | 75,11 |

| a99sb4pew* | Ref_T | Ref_F |
|---|---|---|
| **Pred_T** | 14,84 | 7,71 |
| **Pred_F** | 4,21 | 73,25 |

| Confusion matrix | Ref_T | Ref_F |
|---|---|---|
| **Pred_T** | TP | FP |
| **Pred_F** | FN | TN |

| a99disp• | Ref_T | Ref_F |
|---|---|---|
| **Pred_T** | 14,07 | 7,18 |
| **Pred_F** | 4,98 | 73,77 |

| c22s3p• | Ref_T | Ref_F |
|---|---|---|
| **Pred_T** | 14,02 | 7,16 |
| **Pred_F** | 5,03 | 73,80 |

| a99sbildn4pd• | Ref_T | Ref_F |
|---|---|---|
| **Pred_T** | 14,16 | 7,88 |
| **Pred_F** | 4,88 | 73,07 |

| a14sb3p# | Ref_T | Ref_F |
|---|---|---|
| **Pred_T** | 14,30 | 9,18 |
| **Pred_F** | 4,43 | 72,09 |

| a99sbCufix3p• | Ref_T | Ref_F |
|---|---|---|
| **Pred_T** | 14,17 | 8,99 |
| **Pred_F** | 4,88 | 71,96 |

| a03ws# | Ref_T | Ref_F |
|---|---|---|
| **Pred_T** | 13,98 | 10,82 |
| **Pred_F** | 5,06 | 70,13 |

| c27s3p# | Ref_T | Ref_F |
|---|---|---|
| **Pred_T** | 14,50 | 12,98 |
| **Pred_F** | 4,55 | 67,97 |

**Table S4.** DSSP by residues for references L-shape and U-shape PDBs. C, H, E correspond to Coil, Helix or strand secondary structure, respectively. The RAC2 is showing in the black square.

| Res Seq | Res Id | DSSP U-shape | DSSP L-shape |
|---|---|---|---|
| 50 | TYR | C | C |
| 51 | GLY | C | C |
| 52 | GLN | C | C |
| 53 | SER | E | C |
| 54 | SER | E | C |
| 55 | TYR | E | E |
| 56 | SER | E | E |
| 57 | SER | E | E |
| 58 | TYR | E | E |
| 59 | GLY | C | C |
| 60 | GLN | C | C |
| 61 | SER | E | C |
| 62 | GLN | E | E |
| 63 | ASN | E | E |
| 64 | THR | E | E |
| 65 | GLY | C | E |
